# A porcine germ cell depletion model to investigate the role of germ cells in gonadal development

**DOI:** 10.1101/2025.06.25.661527

**Authors:** Chi-Hun Park, Young-Hee Jeoung, Sai Goutham Reddy Yeddula, JiTao Wang, Bhanu P. Telugu

**Author notes:** Correspondence; 920 E. Campus Dr., 105 ASRC, Columbia, MO 65211.

## Abstract

The molecular mechanisms regulating the germ cell and gonadal somatic cell development are complex and are still poorly understood in non-rodent mammals. The *Nanos* family gene-*NANOS3* is expressed in the migrating primordial germ cells (PGC) and has key roles in germ cell survival in several species. Herein, we investigated the role of *NANOS3* in pigs-an animal model of dual agricultural and biomedical importance. Consistent with prior reports, we found that disruption of *NANOS3* expression did not affect PGC specification. Similarly, migration of PGC to the genital ridge occurs normally, but soon followed by an increase in apoptosis and a loss of undifferentiated state by embryonic (e) day 20. Although *NANOS3^-/-^* gonads initiate development as ovaries or testes, they displayed structural defects. This is especially pronounced in the ovaries after the sex determination period (e35). Notably, we identified defects in pregranulosa and theca cell development at a single-cell resolution, suggesting that germ cells are important for gonadal somatic cell development. This germ cell depletion model therefore offers novel insights into the potential role of germ cells in the emergence of gonadal somatic lineages.

## Introduction

Germ cells are unique cell types that emerge *de novo* during embryonic development and serve as the foundation for genetic transfer to the next generation. In eutherian mammals during the early pre-gastrulation stage of embryonic development, a small subset of primordial germ cells (PGC) arises from the nascent mesoderm called the “primitive streak” in the posterior region of the embryo by upregulation of key “specification” (*PRDM1, PRDM14, SOX17, TFAP2C*) factors (Saitou, 2009). These factors result in suppression of the somatic cell fate, upregulation of the pluripotent factors, and erasure of the imprints, which are reestablished in a sex-specific manner at later stages of gonadal development. The PGC subsequently embark on migration to the genital ridge-the future gonad (Sasaki & Matsui, 2008). In the genital ridges of males, expression of the *SRY* gene in the coelomic epithelial cells results in the upregulation of *SOX9* and a deterministic differentiation to pre-Sertoli cells in the embryonic testes. Conversely, lack of *SRY* expression in females (no Y chromosome) will result in the default pathway, which is the expression of *FOXL2* and differentiation towards the pre-granulosa cells. The pre-Sertoli or pre-granulosa cells in conjunction with the post-migratory PGC organize into primitive sex cords, which remain intact in testes and form the seminiferous tubules or undergo nest breakdown and become primordial follicles in the ovaries. The PGC subsequently undergo differentiation into gonocytes in the testes or enter into meiosis and undergo arrest at the prophase stage of meiosis-I and become a primary oocyte in the resulting ovaries, respectively. Taken together, the development of gonads is a well-orchestrated event and is a result of synchronous and coordinated development of germ cells and their supporting somatic “nurse” cells (Licatalosi, 2016; Saga, 2008).

There are several evolutionarily conserved genes and gene families that play a critical role in germ cell development. One such evolutionarily conserved family is NANOS. The NANOS gene family encode RNA-binding proteins, which serve as translational regulators and are involved in multiple processes of germ cell development including the cell cycle, maintenance of pluripotency, epithelial–mesenchymal transition, and germ cell survival (Haraguchi *et al*, 2003). There are three *Nanos* paralogs, *Nanos1*, *Nanos2*, and *Nanos3* in mice and other eutherian mammals. The Nanos proteins have a typical carboxy-terminal zinc finger motif (CCHC)_2_, which is a highly conserved protein domain that mediates binding with different molecular interaction partners such as the C-terminal RNA-binding domain of Pumilio (Sonoda & Wharton, 1999) as well as with the CCR4–NOT deadenylase complex (Bhandari *et al*, 2014). In mice, Nanos2 binds to the CCR4–NOT complex through CNOT1, whereas Nanos3 does this mainly via direct recruitment of the CCR4-POP2-NOT deadenylase (Suzuki *et al*, 2014). Unlike *Nanos2* and *Nanos3*, which are expressed exclusively in germ cells and have a role in germ cell development (Tsuda *et al*, 2003), mouse *Nanos1* is not expressed in germ cells and is dispensable for germ cell development (Haraguchi *et al*., 2003). The primary function of Nanos2 and Nanos3 in germline cells is to suppress apoptosis by repressing the translation of the pro-apoptotic Bcl-2 Associated X-protein (*BAX*) gene (Suzuki *et al*, 2008). Targeted disruption of *Nanos2* in mice leads to infertility exclusively in the male germline, whereas *Nanos3*-deficient mice exhibit both male and female infertility (Suzuki *et al*, 2007; Tsuda *et al*., 2003). Both *Nanos-2* and *-3* have orthologs in mammals.

Several recent reports from non-rodent studies confirmed their similar conserved and essential role in germ cell development. For example, consistent with the knockout phenotype of *Nanos2*-null mice, *NANOS2*-deficient pigs, goats, and cattle exhibited a loss in male germ cells, but females with deficiencies in these genes remain fertile (Ciccarelli *et al*, 2020; Park *et al*, 2017). A deficiency in *NANOS3* gene manifests as infertility in fetal bovine ovaries (Ideta *et al*, 2016; Mueller *et al*, 2023) and the postnatal pig ovary and testis (Kogasaka *et al*, 2022). Another study has reported that knockdown of *NANOS3* in human embryonic stem cells resulted in a reduction in germ cell numbers during differentiation *in vitro* (Julaton & Reijo Pera, 2011). These findings clearly demonstrate that *NANOS3* is essential for germ cell development in both sexes. Although the loss of *NANOS3* in large animal models leads to germ cell depletion and sterility in the postnatal period (Kogasaka *et al*., 2022), the role of *NANOS3* during PGC migration, colonization and gonadal development, including somatic cell differentiation have not been examined systematically and remain unexplored (Guigon & Magre, 2006; Rios-Rojas *et al*, 2015).

In this study, we generated *NANOS3*-deficient pig fetuses and investigated how the loss of germ cells influences gonadal development and more importantly the gonadal supporting cell differentiation. Extending on previous studies, our results identified the essential role of the *NANOS3* gene in germ cell survival, and the influence of germ cell depletion on gonadal somatic cell differentiation (Kogasaka *et al*., 2022). Our analysis uncovered insights into the transcriptomic features of *NANOS3^-/-^*fetal gonads with cell type-specific developmental defects. These observations provide a basis for future studies on the influence of germ cell depletion on ovarian and testicular development in this and other mammalian species.

## Results

### Functional disruption of *NANOS3* results in the apoptosis of porcine PGC during migration

We sought to investigate whether functional disruption of *NANOS3* leads to apoptosis of PGC during migration. To address this question, we utilized both male and female biallelic *NANOS3^-/-^* embryonic fibroblasts that contained an in-frame premature stop codon in the *NANOS3* gene as nuclear donors in SCNT to generate XX and XY *NANOS3^-/-^* fetuses (Figure 1A). Following a round of SCNT, at e21, the pregnant surrogate was humanely euthanized to recover 6 *NANOS3^-/-^* (4 for XY and 2 for XX) and 4 WT^EGFP^ control XY cloned fetuses (Figure 1B and Figure S1C, D, and Table S3). Additionally, AI was performed with semen from EGFP Tg boar, and 3 age matched WT GFP XX fetuses were recovered for comparison. As expected, loss of *NANOS3* did not result in any overt morphological abnormalities as evident from measurements of the crown-rump length (CRL) and weights of the fetuses, in comparison to their WT counterparts (Figure 1C).

**Figure 1.**
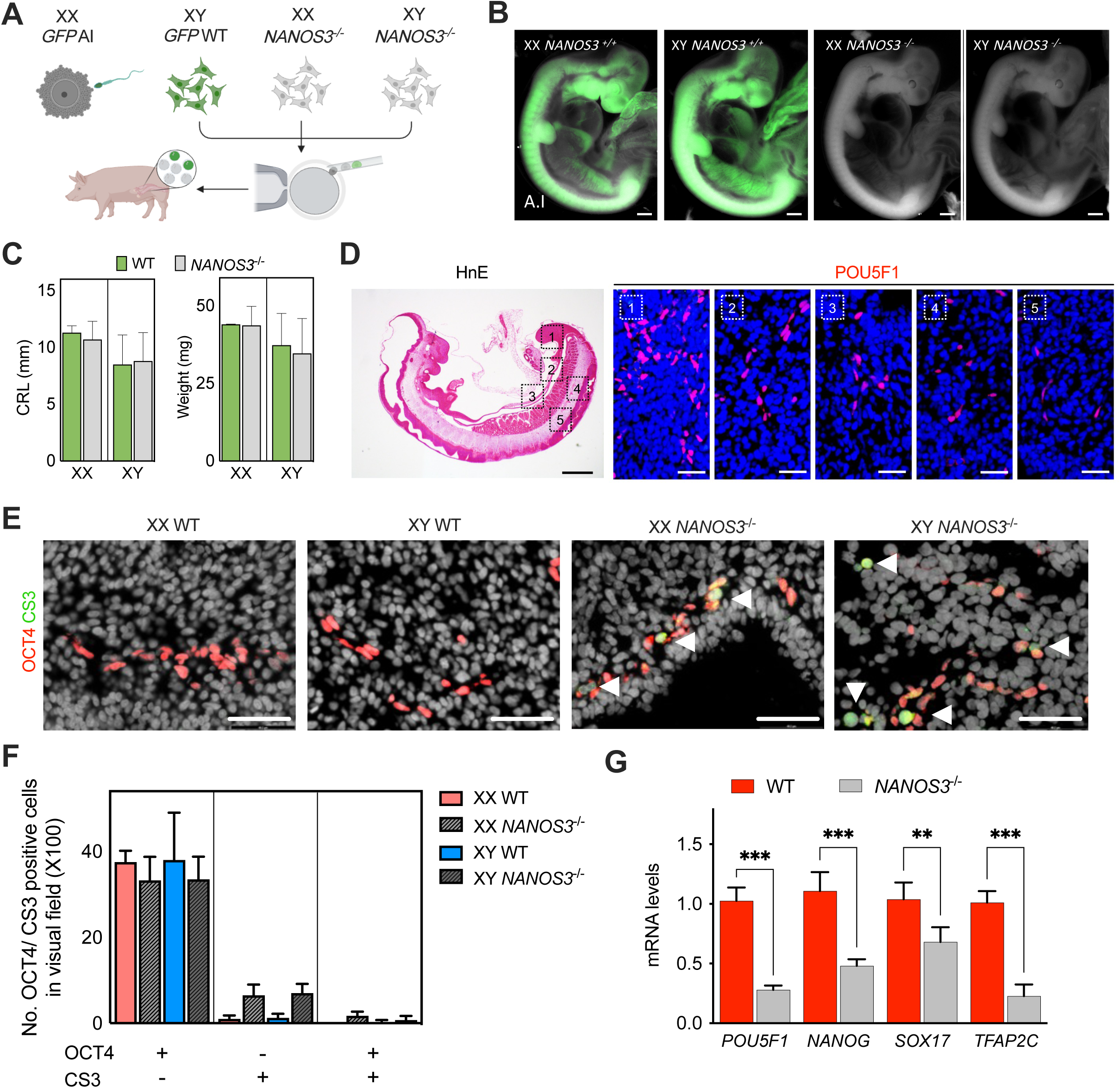
NANOS3 is required for survival of primordial germ cells during migration. (A) Schematic description of the embryo production strategy. (B) Photographs of WT^GFP^ and *NANOS3^−/−^* fetuses followed by SCNT. Bar, 1 mm. (C) Generation of *NANOS3^−/−^* cloned embryos at e21. Crown-rump length (CRL) and body weight of the resulting fetuses. Data are presented as the mean ± S.E. (D) Immunofluorescence staining (bar; 50µm, right) confirmed that the migrating primordial germ cells were POU5F1 positive in the dorsal mesentery zoomed in area from the dotted boxes in the Hematoxylin-eosin (H&E) stained whole fetus (bar; 1mm, left). (E) Double immunostaining with POU5F1 (red) and CS3 (green) to detect apoptotic PGC. The number of CS3+ PGC (arrowheads) was increased in *NANOS3*^−/−^ embryos. Nuclei were counterstained with DAPI (gray). Bar, 50µm. (F) The number of apoptotic PGC (POU5F1+/CS3+) was determined by manually counting cells in serial sections (n=4). (G) Expression levels for representative pluripotent genes. ** P < .01, *** P < .001. Error bars represent means ± S.E.

To assess the impact of *NANOS3* deficiency on migratory PGC, we examined the histological sections for the distribution of cKIT and POU5F1 + (or OCT3/4-a marker for PGC) cells in e21 fetuses (Figure S1E). In the *NANOS3^-/-^* fetuses, we found POU5F1+ PGC along the dorsal mesentery - their migratory route (Figure 1D). This indicates that the PGC specification and migration was not impacted by the disruption of *NANOS3* (Figure S1D)(Kanamori *et al*, 2019). Next, we investigated whether the rate of apoptosis is higher among the migratory PGC in *NANOS3^-/-^* fetuses. To address this, we performed immunostaining for cleaved CASPASE3 (CS3), a marker for apoptosis, in combination with POU5F1 (Figure 1E). We noticed that the proportion of migratory POU5F1+ cells in *NANOS3^-/-^* fetuses (XY: 34.8 ± 7.5; XX: 33.8 ± 5.2) remained comparable to that of the controls (XY: 38 ± 11.0; XX: 37.5 ± 2.6). However, there was an increase in the number of CS3+ apoptotic cells in *NANOS3^-/-^*fetuses (XY: 7.0 ± 1.6; XX: 6.5 ± 1.9) compared to the WT controls (XY: 1.3 ± 0.9; XX: 1.0 ± 0.6; P < 0.05). While CS3+ cells were abundant in the *NANOS3^-/-^*fetuses, only a few cells exhibited colocalized expression of POU5F1 and CS3 (XX, 1.8 and XY, 0.75 on average) (Figure 1F). This reinforces that the population of PGC undergoing apoptosis downregulate POU5F1 expression. To follow-up on these preliminary findings, we performed magnetic activated cell sorting (MACS) enrichment of male WT and *NANOS3^-/-^* fetuses (n=2, each) using SSEA1 antibody-coupled microbeads, followed by co-staining with POU5F1 and CS3 and flow cytometry analysis. There is an approximately 5-fold increase in the number of CS3+ apoptotic cells within the SSEA1+ population of the mutant embryos compared to the WT controls. These results indicate that the loss of *NANOS3* results in a loss of *POU5F1* and pluripotent gene expression followed by an increase in apoptosis (CS3+ cells) (Gkountela *et al*, 2013; Julaton & Reijo Pera, 2011) (Figure1G).

### Loss of germ cells in *NANOS3****^-/-^*** gonads following sex differentiation

Next, we investigated whether the functional inactivation of NANOS3 results in a complete loss of germ cells following the sex determination phase when the bipotential gonads commit to either an ovarian or a testis fate (Parma *et al*, 1999). To address this, we generated XX and XY *NANOS3^-/-^*fetuses via SCNT and compared them to age-matched WT controls at e35 (Figure S2A). Following a round of SCNT, 8 XX and 14 XY fetuses were recovered (Figure S2B). We also recovered developmentally delayed fetuses as assessed by CRL, a common outcome following SCNT. The age-matched “normal” fetuses displayed no differences in weight compared to WT (Figure S2C). The testes exhibited a bean-like appearance with major surface arteries, and were easily distinguishable from the ovaries, which appeared partially twisted and stretched (Coveney *et al*, 2008), indicating that the gonads developed normally into the testis or ovary in a sex-specific manner (Figure 2A). The isolated gonads similarly displayed no differences in the size or weight between the WT and *NANOS3* mutants (Figure 2B).

**Figure 2.**
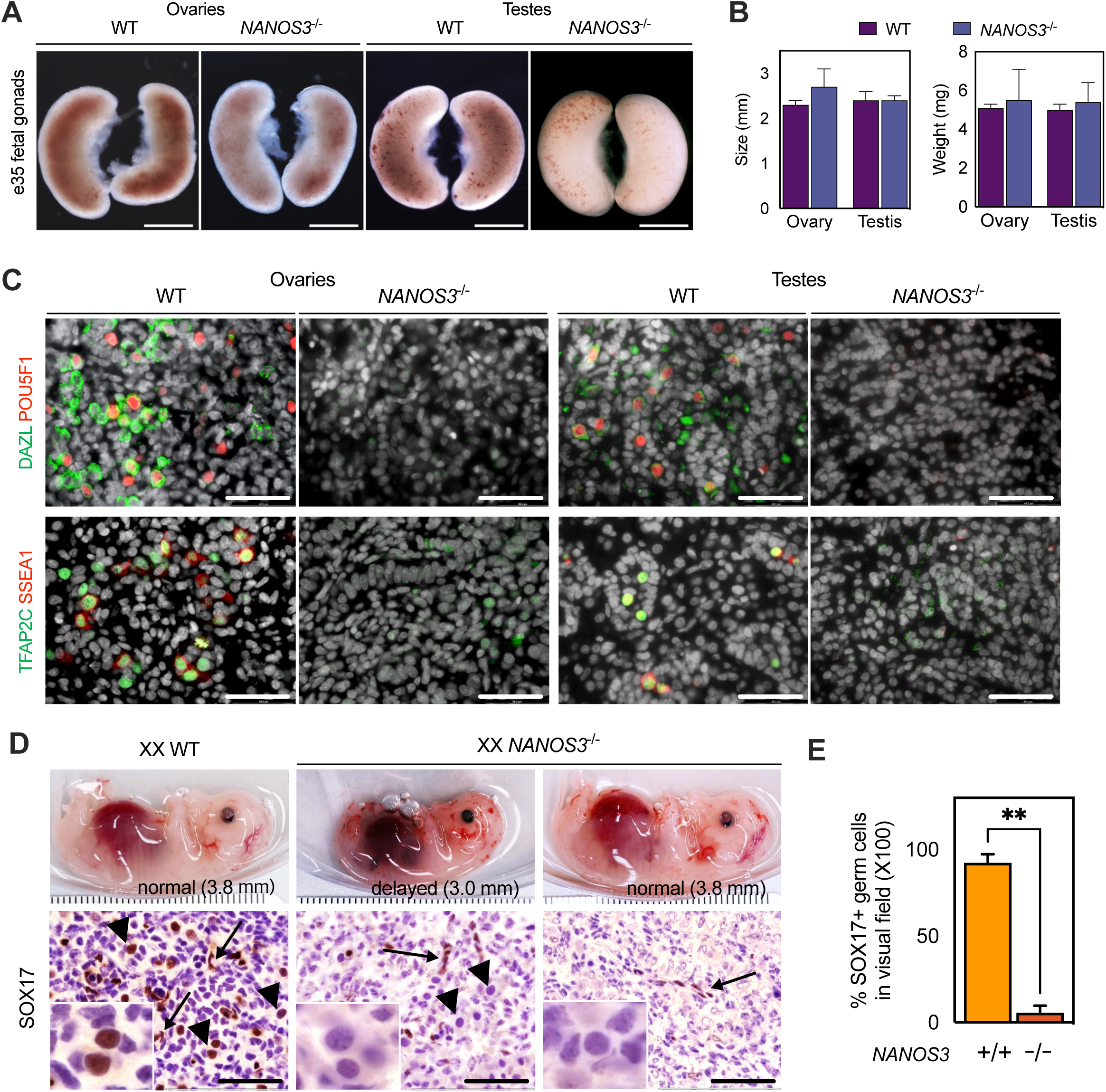
The absence of *NANOS3* causes a complete loss of germ cells following sex determination. (A) Photographs of the WT and *NANOS3^−/−^* gonads at e35. *NANOS3^−/−^* gonads appear to have normal morphology. The testes exhibited a bean-like appearance, along with the arteries, and were easily distinguishable from the ovaries, which appeared partially twisted and stretched. Bar, 1mm (B) Bar graphs illustrate gonad size and weight, as indicated. (C) Immunofluorescence for the early germ cell markers, DAZL (green) POU5F1 (red)-top and TFAP2C (green) SSEA1 (red)-bottom, indicated the absence of germ cells in *NANOS3*^−/−^ ovary and testis. Nuclei counterstained with DAPI (gray) in merged image. Bars, 50 μm. (D) Photographs of WT and delayed/normal mutant fetuses (top) and representative images (bottom) immunostained with SOX17, which marked germ cells (arrowheads, inlet) and endothelial cells (arrow). (E) The total number of SOX17+ germ cells were obtained from counting in slides (n=3). ** P < .01. Error bars represent means ± S.E.

To confirm the germ cell depletion in e35 fetuses (CRL: 38 mm), we investigated for the expression of definitive germ cell-related proteins in the gonads, including the markers for early germ cells, POU5F1, TFAP2C (also known as AP2γ), SSEA1, and DAZL (late germ cells) (Figure 2C). As expected, none of these markers were detected in e35 *NANOS3^-/-^*gonads, confirming the definitive loss of early and late germ cells. To further confirm the presence or loss of PGC, immunohistochemistry with an antibody to SOX17 (with nuclear localization) was performed. The SOX17+ germ cells exhibit large nuclei characteristic of germ cells. Using these criteria, the number of germ cells from each gonad was counted in three independent high-power fields (HPF) (Figure 2D). In e35 WT ovaries, an average of 23 germ cells per HPF (400 total, 5%) were observed with a significant majority (92.3%) showing strong immunoreactivity for the SOX17 (Figure 2E). Interestingly, sporadic SOX17 negative germ cells (1.6%) were identified in two *NANOS3^-/-^* ovaries (7 cells per HPF, 420) from developmentally delayed fetuses (CRL: < 30 mm) (Figure 2D) corresponding to e30 in development. However, no surviving germ cells were found by e35 (CRL: 38 mm). Only endothelial cells were positive for SOX17 (Figure S2D). From these results, it is clear that few germ cells colonized the gonads from e30 fetuses, and by e35, have either undergone apoptosis or lost pluripotency and germ cell identity (differentiation) (Hayashi *et al*, 2004) in both the ovaries and testes.

### Histological architecture of *NANOS3^-/-^* gonads was altered

In the WT ovaries and testes, the primary sex cords (ovigerous nests/cords and testicular cords, respectively) were well organized containing germ cells, and were surrounded by the stromal/interstitial cells (Karl & Capel, 1998) (Figure 3A). However, the ovaries and testes in *NANOS3^-/-^* fetuses were disorganized, and appeared as epithelial monolayers with enlarged cells and abundant finely granular cytoplasm, potentially undergoing degeneration (Figure 3A) (Ideta *et al*., 2016). In the *NANOS3^-/-^* testes, the testicular cords were intact, however, the cords lacked a lumen. We investigated further whether the defects in the cord structure were due to apoptosis or reduced proliferation in the developing gonads, both of which are critical processes involved in cellular homeostasis. We performed immunostaining with the proliferation marker Ki-67 along with DBA (germ cell marker) antibodies (Figure 3B). Interestingly, we identified a significant increase in the number of proliferating cells (Ki-67+) both in the *NANOS3^−/−^* ovaries and testes, compared to the control WT gonads (Figure 3C). We also performed co-immunostaining with CS3 an apoptotic marker, and CDH2-a marker for supporting cells. A key observation was an increase in the number of CDH2+ supporting cells throughout the *NANOS3^−/−^* gonads, compared to the WT counterparts (Figure 3D). Additionally, CS3+ apoptotic cells were significantly higher in the *NANOS3^−/−^*testes and ovaries (5% of the total cells per visual field), compared to the WT controls (<1% of the total cells) (Table S4); however, the apoptosis was restricted to the interstitial/stroma region in mutant gonads. Taken together, the somatic cells proliferate in the absence of *NANOS3* but cannot structurally organize into sex cords in the absence of germ cells, potentially due to missing cell types (germ cells) to rebuild (Gurtner *et al*, 2008).

**Figure 3.**
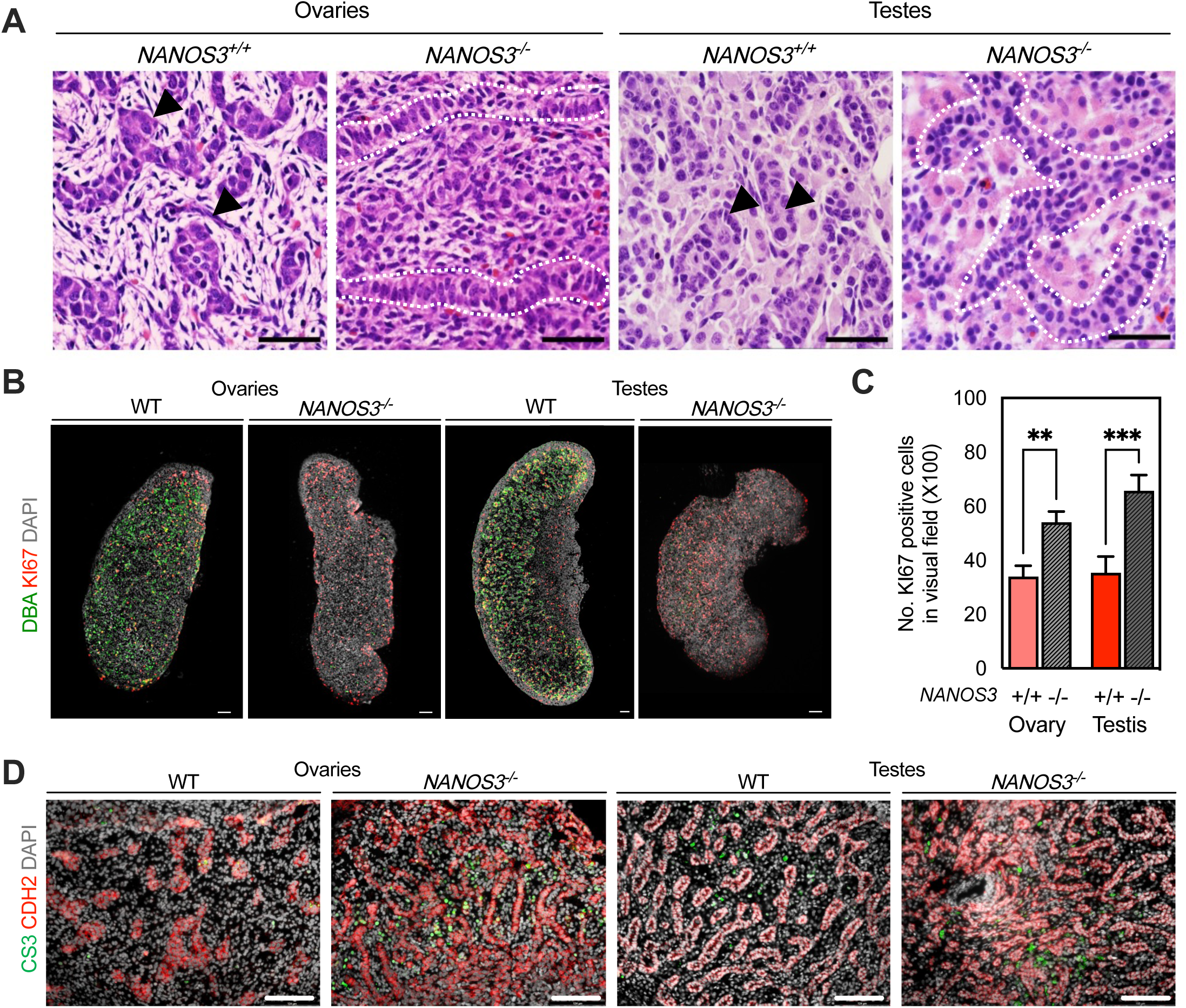
Excessive cell proliferation and apoptosis in *NANOS3^−/−^* gonads. (A) H&E stained histological image of the WT and *NANOS3^−/−^* gonads at e35. Germ cells (arrowheads) along with primary sex cords were formed in WT gonads, whereas the *NANOS3^−/−^* gonads showed loss of germ cells along with disarranged cellular layers in ovaries or with undeveloped tubular structures in testes, which are outlined in dotted white line. Bar 50 µm. (B) Whole mount immunostaining with DBA (green) and Ki67 (red). In the wild type, cells co-expressing DBA (green) and Ki67 were detected in ovaries and testes. DBA expression was not detected in *NANOS3*^−/−^ gonads, whereas *NANOS3*^−/−^ gonads have more Ki67-expressing cells than those of WT control. Bar 100 µm. (C) The total number of Ki67-expressing cells were obtained from counting in slides (n=3). ** P < .01, *** P < .001. Error bars represent means ± S.E. (D) Immunostaining with CS3 (green) and CDH2 (red). Sex cords and unorganized epithelial sheets were detected by immunostaining for CDH2 in all samples. Nuclei were counterstained with DAPI (gray). Bar, 100 μm.

### Single-cell transcriptomic profiling of fetal ovaries and testes of WT and *NANOS3*^-^**^/-^ mutants**

To better characterize the mutant phenotypes in the ovaries and testes at a single-cell resolution, we performed high-throughput snRNA-seq of ovaries and testes after sex determination (e35) from *NANOS3^-/-^* and WT controls by using the 10x Genomics platform (Figure S3A, B). UMAP analysis revealed that ovarian cells from WT (n=4090) and *NANOS3^-/-^* (n=3309) exhibited little overlap, whereas testicular cells from WT (n=5485) and *NANOS3^-/-^*(n=7027) had a greater overlap (Figure 4A). Using the k-means clustering (Figure 4B), the data was partitioned into the following primary gonadal cell lineages based on well-established markers (Uhlen *et al*, 2005): coelomic epithelial cells (**CE**; *GATA4+, ARX+, LHX9+^high^, HHLA2+*), which differentiate into the bipotential early supporting gonadal cells (**ESGC**s; *NR5A1+^high^, LHX9+^high^*,*WT1+^high^, WNT6+^high^*) that give rise to granulosa and Sertoli cells, mesenchymal interstitial/stroma cells (*ARX+, TCF21+, PDGFRA+, COUP-TFII, NR2F2+*), germ cells (**GC**; *DDX4+*), Leydig and theca (*STAR+*) cells, and a relatively small cluster of immune (*CD167+*) cells in the testes and ovaries, respectively (Figure. 4C).

We also evaluated a heatmap of known marker expression across the major cell types (Figure. 4D, E). Among them, CE, interstitial/stromal, and immune cells strongly overlapped, whereas the supporting and interstitial cell populations from WT and mutants from both sexes exhibited minimal overlap within the UMAP space (Figure 4A). In the absence of germ cells in the *NANOS3^-/-^* gonads, we observed distinct changes in WNT6+ ESGCs. Specifically, the ratio of WNT6+ ESGCs in *NANOS3^-/-^* gonads was higher than in WT (19.4% of *NANOS3^-/-^* vs 3% of WT ovaries and 22% of *NANOS3^-/-^*vs 9% of WT testes in population) (Figure. 4F). This is a novel finding not previously reported within these models. The DDX4+ germ cells clustered distantly from the somatic clusters in the WT gonads in both the sexes, which were absent in the *NANOS3^-/-^*gonads (Figure 4B). Based on the expression patterns of the germline-specific genes (Saitou & Miyauchi, 2016), two types of germ cells were identified in the ovaries (Figure 4B). A small fraction (4.2%) of early-stage germ cells (**eGCs**), expressing high levels of pluripotency and early germ cell-specific markers including *POU5F1, NANOG, SALL4, PRDM1* (also called BLIMP1), *PRDM14*, *TFAP2C* (Houmard et al, 2009), *DDX4, DAZL, PIWIL1* and *PIWIL2* (Juliano *et al*, 2011). The second type of germ cells were termed meiotic germ cells (**mGCs**; 11.4%) expressing high levels of the retinoid acid (RA)-induced pre-meiotic genes, *STRA8* and *ZGLP1* (Endo *et al*, 2015; Nagaoka *et al*, 2020), and meiosis initiation genes, *PIWIL4, SYCP1, SYCP2, PAX7,* and *TDRD6* (Garcia-Alonso *et al*, 2022), with concomitant downregulation of early germ cell markers (Figure. 4D). In the testes, 6.3% of the cells were classified as eGC expressing *POU5F1* and *TFAP2C*, and rare RA-responsive meiotic genes including *STRA8* (Figure. 4E). The testicular eGC overlapped with ovarian eGCs, and as expected clustered distantly from mGCs (Figure. 4G). This data further confirms the mitotic and pre-meiotic population of eGCs, and the mGCs undergoing a mitotic-to-meiotic transition (Li *et al*, 2017). Further, we investigated the percentage of germ cells (CDH1+) undergoing mitosis in WT ovaries and testes as shown by colocalization with the Ki-67 (Figure 4H). The analysis confirmed that the total number of double positive (CDH1+/Ki-67+) cells was higher in testes (73%, 36 of 49) than in the ovaries (16%, 14 of 87; Figure 4I). These percentages of actively proliferating germ cells are in agreement with our transcriptome analyses.

### Expression dynamics reveals impaired somatic cell lineages in *NANOS3****^-/-^*** ovaries and testes in the absence of germ cells

To further resolve the observed cell-type composition differences in *NANOS3^-/-^* gonads, we further classified the cells into 16 clusters (hereafter abbreviated as “C”) in ovaries, and 19 C in testes using the known markers from previous studies (Garcia-Alonso *et al*., 2022) and based on DEGs (Figure 5A and 5B; Table S5). The molecular phenotype of these various clusters was shown in Table S4. Salient observations from these clustering data for WT and mutant gonads are listed below.

#### Coelomic epithelial (CE) cells

In the case of ovaries, the CE cells (C2, C10, C11) were defined by their expression of *ERBB4, MAMDC2, SBSPON, SOX6, WNT5A*, and *HHLA2,* which could be subdivided into two clusters: a mitotic (C2) and non-dividing (C10, C11) epithelium. The mitotic CE is enriched for proliferation genes such as *TOP2A, RACGAP1, PRC1, BIRC5* (Gao *et al*, 2021; Morris *et al*, 2022; Yang *et al*, 2018), and are likely the primary source of gonadal somatic cells in ovarian development (Figure 5). The ratio of mitotic and non-dividing CE cells in the *NANOS3^-/-^*ovary (10.7% and 9.3%) was comparable to the WT control (11.9% and 8.8%, respectively). In the testis, only 2% of the cells are CE (C18). This indicates that the expansion of proliferating CEs occurs in the ovaries and are distinct from the tunica albuginea somatic cells (Mork *et al*, 2012; Rastetter *et al*, 2014) in the testis expressing *GATA2, KRT19, UPK3B*, which were defined as gonadal ridge epithelial-like cells (Hummitzsch *et al*, 2013).

#### Pregranulosa and Sertoli cells

The ESGC (C1, C3, and C6) constitute the largest proportion of cells in both the WT (32.8%) and mutant (47.8%) ovaries, and express factors such as *RSPO1, KDR, CDH2, KITLG, PTPRN2,* and *LHCGR*, illustrating their identity as gonadal early supporting progenitors. Of note, there were two distinct supporting cell populations in the ovaries. The first population is WNT6+ (C5 and C12) cells, which make up for a significant majority of cells in the mutant ovaries (24.5%) and shared partial similarity with ESGCs (Figure 4, Figure S1E). This observation is suggestive of an earlier induction of the molecular networks underlying the initial commitment of ESGCs to the granulosa fate, termed the “transitional” cells. The second population is LGR5+ (C15) cortical preGCs, which are present exclusively in the WT ovaries (4.8%), and are distinguished by their expression of steroidogenic enzymes (*HSD17B3, NR5A2, INHBA*) and their regulators (*PTPN5, FSHR,* and *KCNK13*) (Gustin *et al*, 2016; Wamaitha *et al*, 2023). In addition, DEGs in the WNT6+ and LGR5+ cells showed enrichment for GO terms related to the WNT signal pathway, ubiquitin mediated proteolysis, growth hormone synthesis, FOXO-mediated transcription among top-enriched gene sets (Table S5). In the testes, the bipotent cells (C3) possess a *PDGFRA*+ interstitial and supporting *SOX9*+ cell identity which are distinct from the ovaries. As expected, the largest population in testes was SOX9+ Sertoli cells, which shared markers with their counterparts-preGCs in the ovaries, including, *LRG5, HSD17B3, and CDH2* (Figure 5E). Contrary to the ovarian phenotype, there were minimal differences in the Sertoli cell population between the WT (36%) and mutant testes (30%).

#### Theca and Leydig cells

In the ovaries, fewer theca cells were observed in *NANOS3^-/-^* ovaries (0.2%) compared to the controls (1.4%). The *PDGFRA+* and *NR2F2+* (C4, C7, C9, C13) interstitial stromal progenitors can be further categorized into the steroidogenic (C4 and C7) and fibroblast-like (C9, C13) populations. This includes the C4 cluster, which has a small fraction of STAR+ theca cells (Fan *et al*, 2019) enriched for steroidogenic genes (e.g., *CYP11A1, CYP17A1, LHCGR, INSL3, FDX1, LDLR, GLDN*). Similar to theca cells, STAR+ Leydig (C14) cells were readily identifiable in WT and mutant testes, but the proportion was not different between them (6.0% and 8.4%, respectively) (Figure 5E). Among the testicular somatic cells, the ratio of *SOX9+* (C10) supporting cells and *PDGFRA+* (C11) interstitial proliferating cells (e.g., *TOP2A*) in the mutant gonads were two times higher than that in the WT (6.7% Vs 3.6% and 7.4% Vs 3.3%) respectively consistent with the previous results (Figure 3B). From these results, we conclude that germ cells contribute to the differentiation of somatic lineage (preGC and theca) in the ovary. However, despite some changes in cellular composition, the differentiation of Sertoli and Leydig cells in the mutant testes remain unaffected in the absence of germ cells. These observations point to a unique sexually divergent regulatory mechanism underlying male and female gonadal supporting cell development, highlighting a profound influence of germ cells on ovarian cell lineage specialization.

## Discussion

Germ cells have a unique lifecycle including-*de novo* specification from the incipient mesoderm, an arduous and perilous journey to their future home – the genital ridge, cooperative organization with the EGCCs into sex cords, and differentiation and development into primary oocytes or gonocytes, in ovaries and testes respectively. However, the underlying mechanisms for germ cell and supporting gonadal somatic cell development, and how they influence each other in the maturation and higher order organization into primitive sex cords remain yet to be fully elucidated in non-rodent species mammals. Historically, while the livestock species, specifically the domestic pig has been considered as an ideal alternative, the lack of genetic engineering tools has hindered their use as models for mechanistic studies. The recent development of genome editors, including the CRISPR/Cas has proven to be a gamechanger for the field. The editors can introduce site-specific breaks in the DNA, which can be leveraged for generating transgenic pigs with precision. Coupled with advances in sequencing at single cell resolution and improvements in bioanalytical tools, it is now possible to use these models to either fill-in the gaps emanating from research in rodents or serve as an alternative. In this study, taking advantage of the progress in genetic engineering, we set out to investigate the effects of germ cell depletion in early gonadal development especially in the context of supporting somatic cell development. This study demonstrated that disrupted *NANOS3* expression led to the loss of germ cells postmigration, defects in the somatic cells especially in the ovary, and overall disorganization of sex cords in both gonads. Our snRNA seq data provided novel evidence that germ cells are essential for gonadal somatic differentiation during ovarian development.

### Investigating the protective role of NANOS3 in fetal germ cells in pigs

Apoptosis is a crucial mechanism underlying many genetic forms (including nanos genes) of germ cell loss in mice and other animal models (Aitken *et al*, 2011). We observed that normal numbers of migrating PGC were present in the *NANOS3* deficient concepti at e21. However, there was an increased rate of apoptosis and decreased expression of pluripotency markers compared to WT (Figure 1). Thus, it is clear that disrupted *NANOS3* expression did not interfere with PGC specification and proliferation, but it became indispensable for germ cell viability during migration via suppression of apoptosis (Tsuda *et al*., 2003). Moreover, our results support the notion that *NANOS3* expression may maintain an undifferentiated state of early fetal germ cells (Julaton & Reijo Pera, 2011; Wang & Lin, 2004; Yamaji *et al*, 2010). Although our results are suggestive and only observed at the gene expression level, this observation implies that NANOS3 may be involved in regulating POU5F1 expression (Gkountela *et al*., 2013; Julaton & Reijo Pera, 2011). Thus, further investigation is warranted to probe the possible role of NANOS3 in self-renewal and maintenance of early pig germ cells.

We further focused on addressing the developmental defects in the mutant gonads after the sex determining period (e35). Consistent with prior studies, our results showed that *NANOS3* deficiency leads to a progressive and complete loss of germ cells in both sexes and continues up until after sex differentiation (Ideta *et al*., 2016; Mueller *et al*., 2023; Tsuda *et al*., 2003). Intriguingly, we found that a small proportion of *NANOS3*-depleted germ cells survived and colonized the genital ridges, as previously described (Tsuda *et al*., 2003). Although the BAX-dependent or independent apoptosis is associated with germ cell loss in such models, it remains unclear as to the exact mechanisms of germ cell loss and the fates of a few germ cells that survive after reaching the genital ridge (Suzuki *et al*., 2008). It is possible that a few surviving germ cells will undergo apoptosis at a later stage or potentially lose pluripotency and differentiate into other lineages (Hayashi *et al*., 2004).

### Impact of germ cell loss on the differentiation of supporting cell lineage in fetal gonads

The influence of germ cells on somatic cell differentiation in the ovarian development has been reported in invertebrate species (Slanchev *et al*, 2005) but has not been investigated and/or reported in mammals. Reports from *NANOS2* or *NANOS3* deficient pigs in which germ cells have been depleted have grossly normal testis at birth; however, exhibit severe hypoplasia in postnatal periods due to lack of spermatogenesis. In these animals, there is evidence of intact seminiferous tubules with functional Sertoli and Leydig cells (Ideta *et al*., 2016; Mueller *et al*., 2023; Park *et al*., 2017). In contrast, the ovaries exhibited defects in follicular structures, with few, if any, true follicles. These observations support the notion that the germ-soma interaction is required for ovarian differentiation and folliculogenesis in the postnatal period (Fukuda *et al*, 2021). In the present study, *NANOS3^-/-^* (e35) ovaries and testes exhibited defects in the cords, with testis showing a lack of lumen, and ovaries containing a sheet of epithelial cells (Ideta *et al*., 2016). These structural alterations suggest that the intercellular interactions mediated by cell-cell contacts and/or paracrine factors may play a pivotal role in the organization of complex gonadal structures following CE ingression (Guigon & Magre, 2006; Nicholas *et al*, 2010).

Our single nuclear RNA-Seq data highlighted the distinct transcriptomic features of the major gonadal populations including germ cells, interstitial/stromal (Leydig/theca), gonadal progenitors, and supporting (granulosa/Sertoli) cell lineages. Several observations are worth mentioning: first, our data indicated that testicular and ovarian cells shared cell type specific markers, despite having distinct developmental pathways. This observation suggests that they retain similar function and characteristics to their roles in the reproductive system through a parallel developmental path. Second, LGR5+ preGC cells were low in *NANOS3^-/-^* ovaries, and instead WNT6+ transitional cells harboring both supporting and stromal identity were enriched in *NANOS3^-/-^* ovaries. In addition, STAR+ theca cells were found in WT ovaries but are lower in mutants, likely suggesting that signals from pregranulosa cells may regulate their differentiation and development (Schmahl *et al*, 2008). Although the transcriptome feature was similar between WT and mutant ovaries, the disorganized cord structure may be a consequence of defects in somatic cells downstream of the germ cell loss in *NANOS3^-/-^* gonads. Although, the degree to which these mechanisms mirror the differentiation of granulosa cells in ovaries—a process central to the formation of follicles—is still being explored. Evidence from this study suggests a role for germ cells in directing the fate of supporting progenitor cells in developing ovaries after sex determination. Intriguingly, such compositional changes in their testicular counterparts, i.e., the Sertoli and Leydig cells, have not been observed in mutant testes. The observed female biased defects reflect that the mechanism and factor(s) involved in cell fate commitment are differentially controlled between the sexes at the sex-determining stage. As to possible mechanisms underlying the sexual dimorphism, differences in the cell cycle stages in fetal germ cells between the sexes may be the underlying cause, as has been shown in re-aggregation experiments. The disruption of somatic cell differentiation observed in our *NANOS3* mutant ovaries is a yet undescribed phenotype and the causative phenomenon still needs to be clarified. Going further, elucidation of mechanisms and factors involved in specification and maintenance of female supporting cell fate may provide novel insights into sex differentiation. Because the ovarian cord formation is conserved in mammalian germ cell development (Pepling *et al*, 1999), the identification of genetic components and molecular mechanisms of fetal stage germ and somatic cell structures may better translate to understanding human gonadal development.

### Conclusions

The cell fate commitment of germ cells and gonadal somatic cells in early development are incompletely characterized and, still, only poorly understood. This study demonstrated that disrupted *NANOS3* expression led to the loss of germ cells postmigration, defects in somatic cell development, and disorganization of sex cords in the developing gonads. Our snRNA seq data provided novel evidence that germ cells are essential for gonadal somatic differentiation during ovarian development. Our data from a germ cell depletion pig model may potentially be used for further elucidating the influence of germ cells on the somatic differentiation during gonadal development. While a strong effort was made to characterize the developmental defects due to *NANOS3* deficiency, our experiments also allowed us to provide comprehensive single-cell transcriptomics resources for gonadal development thereby enhancing our knowledge.

## Materials and methods

All chemicals were purchased from Sigma-Aldrich (St. Louis, MO) unless stated otherwise. All experiments involving live animals were performed as per the approved guidelines of the University of Missouri, Institutional Animal Care and Use Committee protocol # 45081.

### Generation of homozygous *NANOS3*-null and age-matched WT fetuses

To generate *NANOS3* null (*NANOS3^-/-^*) XX and XY fetuses, we utilized the sequence validated biallelic *NANOS3^-/-^*embryonic fibroblasts that contained an in-frame premature stop codon in *NANOS3* as nuclear donors for somatic cell nuclear transfer (SCNT) (Figure S1A) (Park *et al*, 2022). The introduction of the stop codon results in the production of a truncated NANOS3 protein lacking the indispensable zinc finger-containing domains, rendering the protein non-functional (Figure S1B). To generate age matched wild type (WT) control embryos, we utilized either the fetal fibroblasts for SCNT, or artificial insemination (AI) of a gilt in standing estrus with semen from the *EGFP* transgenic (Tg) line (NSRRC Strain # 0016). Timed euthanasia of the surrogates following embryo transfer or AI was performed to recover *NANOS3^-/-^* or WT fetuses for phenotypic analysis. For SCNT, briefly, cumulus-oocyte-complexes (COC) were purchased from a commercial supplier (De Soto Biosciences, Seymour, TN, USA). Following *in vitro* maturation for 40-44 h, the cumulus cells were removed from the oocytes by gentle pipetting in a 0.1% (w/v) hyaluronidase solution. *NANOS3^-/-^*or WT embryonic fibroblasts were synchronized to the G1/G0 phase by serum deprivation (DMEM with 0.1% FBS) for 96 h. The oocytes were enucleated by aspirating the polar body and the MII metaphase plates with a micropipette (Humagen, Charlottesville, VA, USA) in 0.1% DPBS supplemented with 5 μg/mL of cytochalasin B. After enucleation, one donor cell was placed into the perivitelline space of an enucleated oocyte. The cell– oocyte couplets were fused by applying two direct current (DC) pulses (1 s interval) of 2.0 kV/cm for 30 μs using an ECM 2001 Electroporation System (BTX, Holliston, MA). After fusion, the reconstructed oocytes were activated by a DC pulse of 1.0 kV/cm for 60 μs, followed by post-activation in 2 mM 6-dimethylaminopurine for 3 h (Park *et al*, 2021). After overnight culture in MU4 medium with a histone deacetylase inhibitor Scriptaid (0.5 μM), the reconstructed zygotes were cultured in MU4 in low oxygen (5% O_2_ and 5% CO_2_ in 90% N_2_) for embryo transfer (Chen *et al*, 2021; Park *et al*., 2021). The cloned embryos were surgically transferred into the oviduct of estrus synchronized recipients on the first day of standing estrus. Timed euthanasia of the surrogates following embryo transfer or AI was performed to recover *NANOS3^-/-^* or WT fetuses for phenotypic analysis.

### Genotyping and sexing analysis

Extraembryonic membranes from the fetuses were utilized for isolating DNA by using a DNA isolation micro-Kit (Norgen) according to the manufacturer’s instructions. Genotyping and sexing were performed via PCR by using either KOD Hot start mastermix (Novagen) or Bioline Taq under the following conditions: denaturation and polymerase activation step of 95°C /3 min, 35 cycles of 95°C /20 s, 60°C /10 s, 70°C /10 s, and the final extension step of 70°C /5 min. The sex of WT controls was determined by genotyping by using the pig Y chromosome, *SRY*, as previously described (Park *et al*, 2012). The primers used were: 5’-*CTGGGATGCAAGTGGAAAAT*-3′ (forward) and 5′-*GGCTTTCTGTTCCTGAGCAC*-3′ (reverse); the PCR program used was 5 min 95°C, 34 cycles 95°C /1 min, 60°C /30 s, 72°C / 2 min, 72°C /10 min, and the PCR products were resolved on a 1.5% agarose gel. The PCR products were purified using NucleoSpin gel and PCR clean-up kit (Machery Nagel, Bethlehem, PA), and were subjected to Sanger sequencing (using the forward PCR primer; Table S1).

### Histology and immunofluorescence

At embryonic day (e) 21 and 35, whole concepti and gonads from the fetuses were recovered, respectively, and were fixed in 4% paraformaldehyde (PFA) in PBS at 4°C overnight followed by long-term storage in 70% ethanol at 4°C. Conceptus or fetal gonads were dehydrated, paraffin embedded and sectioned for immunohistological analysis. For immunofluorescence, heat-induced epitope retrieval was conducted in sodium citrate buffer (10 mM sodium citrate, 0.05% Tween-20, pH 6) at 95°C for 20 min. Subsequently, the sections were permeabilized with 1% Triton X-100/PBS for 15 min and blocked with superblock (Fisher Scientific) or 5% BSA for 1 h prior to antibody incubation. Primary and secondary antibodies used for immunofluorescence are listed in Table S2. Incubation with the primary antibodies was performed at 4°C overnight. The next day, cells were washed and incubated with fluorescent (TRITC, Cy3 Alexa fluorophore 488, 555, 568 and/or 647; eBioscience, Abcam, Thermo Fisher Scientific)-conjugated secondary antibody for 1 h at room temperature (RT). Slides were then washed three times with 0.1% Tween-20/PBS. The nuclei were counterstained with 5 mg /L DAPI (Fisher Scientific) during the final washing series. Slides were mounted with diamond antifade mounting solution and stored at −20°C until imaging. Bright-field images of the embryos and the gonads were acquired by using Leica DFC 500 and DFC 7000T cameras. Fluorescence images were photographed by using a Leica DM 4000B microscope using the Leica Application Suite Advanced Fluorescence software (Leica Microsystems, Wetzlar, Germany).

### Quantitative analysis of apoptotic population

The apoptotic population within the gonads was examined by semiquantitative evaluation of the expression of cleaved-CASPASE3 (CS3) and POU5F1, which are apoptotic and germ cell markers, respectively. Ki67 immunohistochemistry was performed to determine proliferating cells in the ovary and testis. The apoptosis/ proliferation indexes were calculated by dividing the number of CS3 positive (+) or Ki67+ cells by the total number of cells in randomly selected fields. The average values of cells in the knockout gonads were compared to the WT control, and statistically analyzed.

### Flow cytometry analysis

Whole concepti were dissociated by using gentleMACS™ Dissociator (Miltenyi Biotec, Cat#130-093-235) with 0.25% trypsin/EDTA at 37°C for 5–15 min and bound with Anti-CD15 (SSEA1) MicroBeads (Biolegend) and subjected to FACS Frotessa cytometry (BioRad). FACS data was analyzed by Flowjo software.

### Single-Nuclei Isolation, sequencing using 10x Genomics protocol and data analysis

Whole gonads were washed in ice-cold 100% MeOH and stored at - 80°C for single-nuclei extraction. Single nuclei were isolated from around 20-30 mg of three pooled gonads from each group by using a Singulator 100 machine (S2Genomics, Livermore, CA), following the manufacturer’s instructions. After isolation, nuclei preparations were centrifuged at 500 g for 5 min and resuspended in 2 mL of a buffer supplemented with 1 U/µL of RNAse inhibitor (Fisher). Nuclei stained with Trypan blue were inspected under an inverted microscope manually. Nuclei concentration was estimated by DAPI staining on a Countess II FL automated cell counter ⩾ 2 times for each sample (Figure S3 <u>A,B</u> (Becht *et al*, 2018; Raudvere *et al*, 2019; Zhang *et al*, 2023). Libraries were constructed by following the manufacturer’s protocol with reagents supplied in the 10x Genomics Chromium Next Gel Bead-in-Emulsion (GEMs) Single Cell 3′ Kit. Single nuclear sequencing was performed by the Informatics Research Core Facility, University of Missouri, Columbia. The data was pre-processed with the CellRanger software (v7.0.0), including demultiplexing, fastq file generation, read alignment, and UMI counting (Zheng *et al*, 2017). The WT and *NANOS3*-/-datasets were integrated for each sex separately. The obtained data were analyzed and visualized by using the 10x Genomics Loupe Browser v6.5.0 software (10x Genomics Inc.). We analyzed the expression of known markers for the expected cell types and assigned identities to the clusters. Differentially expressed genes (DEG) between WT and *NANOS3^-/-^*were used as input for gene ontology enrichment analysis by web-based g:Profiler (Raudvere *et al*, 2019).

### Data Availability

A total of 4 snRNA-seq data sets generated for this article can be accessed via NCBI Sequence Read Archive (SRA) with the accession number PRJNA1161363.

### Statistical analysis

The statistical analysis was performed by using GraphPad Prism software 9.0 (GraphPad Software, La Jolla, CA, USA). The data was expressed as mean ± SD. For normally distributed data, t-test was used for comparison between 2 independent samples, and one-way analysis of variance (ANOVA) was used for comparisons of multi-samples. A P-value of <0.05 was considered as statistically significant. The number of replicates in each experimental setting and statistical significance are shown in each figure legend.

## Acknowledgments

This research was supported by investigator start-up funds and by Genus PlC, a publicly listed animal genetics company. The funder was not involved in the study design, collection, analysis, interpretation of data, the writing of this article or the decision to submit it for publication.

## Author contributions

C.P. and B.P.T. conceived the project and designed the experiments; C.P., Y.J., S.G.Y., and J.W. established and performed characterization of the phenotype; C.P., and S.G.Y. generated the libraries and transcriptomic analysis; C.P. wrote the initial draft; B.P.T. revised the manuscript based on the input from all authors. All authors approved the final draft for submission.

